# MICROENDOSCOPIC CALCIUM IMAGING IN SUPPLEMENTARY MOTOR AREA AND PRIMARY MOTOR CORTEX OF RHESUS MACAQUES AT REST AND DURING ARM MOVEMENT

**DOI:** 10.1101/2024.06.20.599918

**Authors:** Anne-Caroline Martel, Damien Pittard, Annaelle Devergnas, Benjamin Risk, Jonathan J. Nassi, Waylin Yu, Joshua D. Downer, Thomas Wichmann, Adriana Galvan

## Abstract

The study of motor cortices in non-human primates is relevant to our understanding of human motor control, both in healthy conditions and in movement disorders. Calcium imaging and miniature microscopes allow the study of multiple genetically identified neurons with excellent spatial resolution. We used this method to examine activity patterns of projection neurons in deep layers of the supplementary motor (SMA) and primary motor areas (M1) in four rhesus macaques. We implanted gradient index lenses and expressed GCaMP6f to image calcium transients while the animals were at rest or engaged in an arm reaching task. We tracked the activity of SMA and M1 neurons across conditions, examined cell pairs for synchronous activity, and assessed whether SMA and M1 neuronal activation followed specific sequential activation patterns. We demonstrate the value of *in vivo* calcium imaging for studying patterns of activity in groups of corticofugal neurons in SMA and M1.

**HIGHLIGHTS:**

- Use of one-photon miniature microscopes and microendoscopic calcium imaging to study the activity of cortical projection neurons in the supplementary motor area (SMA) and primary motor cortex (M1) of rhesus macaques at rest or performing simple arm reaches.
- Calcium transients were related to arm reaches and showed directional sensitivity in a proportion of cells in SMA and M1.
- Subsets of cell pairs showed coactivation in SMA and M1 during rest and reaching tasks. The strength of coactivity was not related to the distance between cells.
- SMA and M1 neurons displayed sequential activation patterns.
- We demonstrated that microendoscopic calcium imaging can be used to assess dynamic activity within genetically identified cell populations in deep layers of SMA and M1.

## INTRODUCTION

Cortical motor regions have undergone significant evolutionary expansion, specialization, and re-organization ^1^, therefore non-human primates (NHPs) are often used to model motor control in humans under healthy conditions, and during movement disorders such as Parkinson’s disease. In primates, the motor cortices include the primary motor cortex (M1) and the supplementary motor area (SMA) ^2,3^, which are components of larger frontal networks that also involve portions of the basal ganglia and the ventral anterior and ventral lateral thalamus ^4–7^. Neurons in M1 and SMA are involved in movement planning and execution ^2,8^.

The majority of studies on cortical neuronal activity in awake NHPs have used extracellular electrophysiological recordings. Methods such as one-photon fluorescent miniature microscope (miniscope) imaging of genetically-encoded calcium indicators offer an alternative approach to monitor neuronal activity, and have recently been used to study calcium transients in individual cells in the premotor and visual cortices of macaques ^9–11^.

When combined with microendoscopic gradient index (GRIN) lenses, miniscopes enable optical access to non-superficial imaging regions, and allow prolonged monitoring of multiple individual neurons in the brain region of interest, providing detailed understanding of spatial relationships among cells. Calcium indicators, such as GCaMP, can be genetically directed to be expressed in selective cell populations, and the resulting calcium activities of identified neurons can be followed throughout the course of an experiment, enabling tracking of the same cells across behavioral conditions (e.g., from rest to active movement).

We used microendoscopic calcium imaging to examine the activity of neuronal ensembles in SMA and M1 in four rhesus macaques while the monkeys were at rest or performing simple arm reaches. Following expression of the genetically encoded calcium indicator GCaMP6f in projection neurons of deep cortical layers of SMA and M1, we tracked the activity of these neurons across conditions, examined cell pairs for synchronous activity or inactivity, and assessed whether neuronal activation followed specific sequential activation patterns. Our results demonstrate that calcium imaging captures dynamic activity patterns in SMA and M1 neurons and establish a basis for future research on how SMA and M1 activities change during the development of parkinsonism and other disorders of movement in NHPs.

## RESULTS

We studied the neuronal activity in SMA and M1 in four macaques using calcium imaging. We used AAVs to express GCaMP6f under control of the Thy1 promoter to target preferentially projection neurons ^12^ in deep cortical layers. Surgical preparations to chronically implant microendoscopic GRIN lenses for optical access to motor cortices and to ensure a stable long-lasting cranial chamber were performed as described in Bollimunta et al. (2021). Out of four subjects, we successfully performed calcium imaging in three SMA sites (monkeys Q, U and F) and two M1 sites (monkeys V and U).

### General description of calcium activity

Calcium imaging was done while the animals were sitting in a primate chair and making occasional spontaneous movements (‘spontaneous’ state), or while they were engaged in a reaching task (monkeys Q and U).

Individual cells and their calcium dynamics were identified using the Constrained Non-negative Matrix Factorization for microendoscopic data (CNMFe) algorithm and validated, based on their activity pattern and morphology (figure 1 A-C; see table S1 for parameters used). The number of identified GCaMP6f-expressing cells varied across imaging sites and sessions. In the SMA, we identified 129, 61 and 63 cells in monkeys Q, F and U, respectively. In M1, we identified 37 and 23 cells in monkeys U and V, respectively. (table S2). We could track a small portion of cells (18%) across up to three recording sessions.

**Figure 1:**
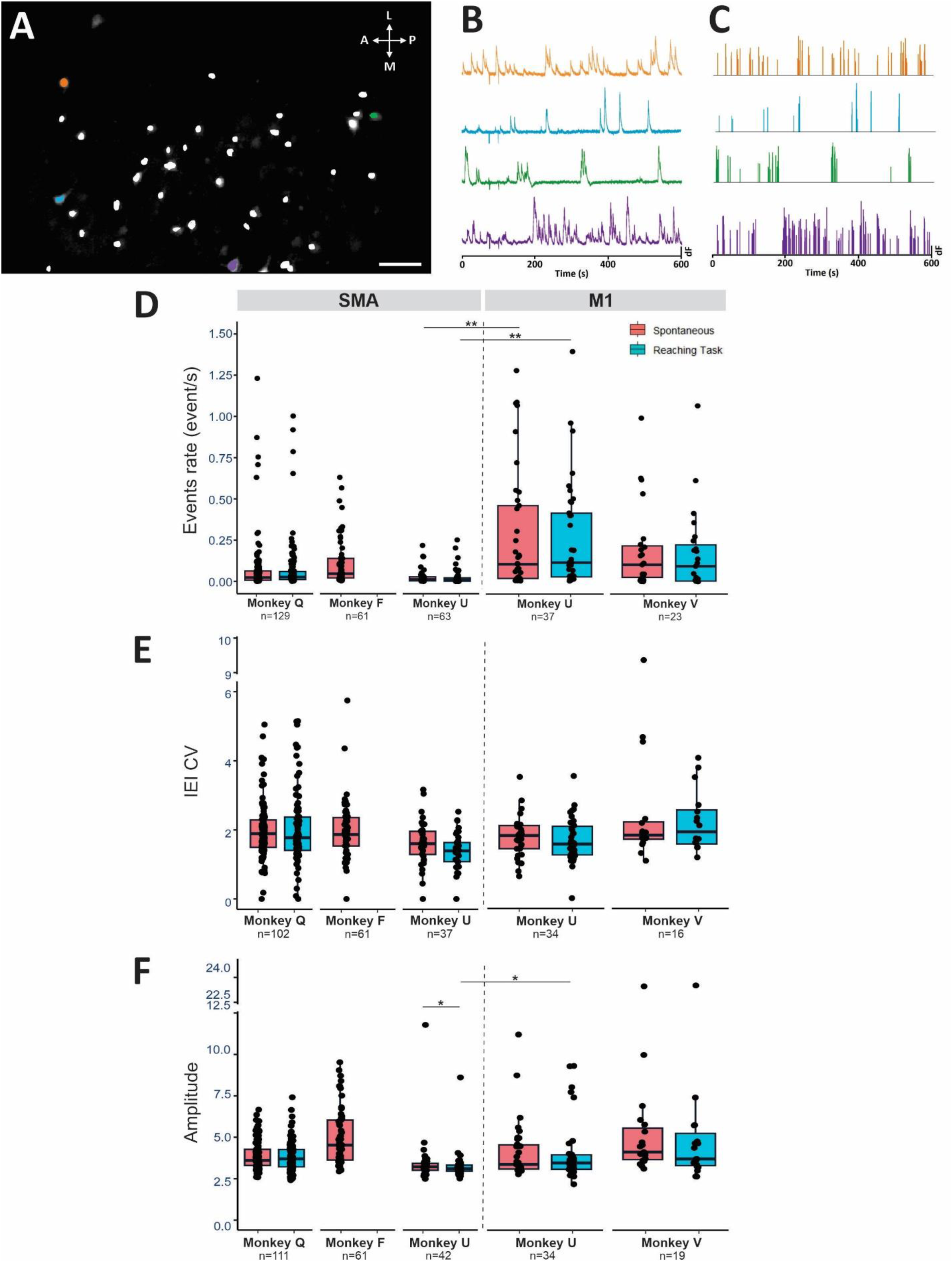
Steps of analysis and general description of cell activity. **A:** Maximal intensity of GCaMP6f fluorescence in the SMA during a single example session (monkey Q), overlapped on the cell map (white) extracted using CNMF-E. Orange, blue, green, and purple symbols mark the example cells shown in B and C. Scale bar: 100 µm. **B:** Raw traces of calcium transients (dF, peak-normalized) during spontaneous activity. **C:** Calcium events deconvolved from the calcium traces in B. **D, E, F:** Box plots summarizing the event rate (events/s), inter-event interval coefficient of variation (IEI CV) and amplitude of events detected across all sessions in the SMA (left) and M1 (right) in the spontaneous condition (red) and during the arm reaching task (blue). The horizontal bar in each box indicates the median, each circle indicates the median value for a cell. * p < 0.05, ** p <0.001; Wilcoxon signed rank tests corrected for multiple comparisons using the false discovery rate (FDR).

Calcium events were extracted from the raw calcium traces using the OASIS deconvolution method (see Methods). A comparison of calcium event kinetics revealed region- and task-dependent responses in SMA and M1 neurons. The calcium event rates and the coefficient of variation of the inter-event intervals (IEI CV) did not differ between the spontaneous and reaching task conditions in either cortical region (figure 1D-E). In the SMA, the amplitude of the events was significantly higher in the spontaneous state, compared to the reaching task recordings in Monkey U (p=0.032; Wilcoxon signed rank test, corrected for multiple comparisons). In Monkey U, we found that the event rate was higher in M1 than in SMA, both during the spontaneous and reaching task conditions (p<0.001 for both, Wilcoxon rank sum test). Similarly, the amplitude of the events was higher for the task in M1, compared to SMA (p=0.03, Wilcoxon rank sum test, figure 1D-F).

### Calcium traces in SMA and M1 show changes in activity in relation to an arm reaching task

We measured the calcium activity of SMA and M1 neurons in relation to behavioral events in an arm reaching task. Monkey Q performed a simple one-target task in which a circle was randomly presented at the right, left or center of a touchscreen. Touching the target within 3 seconds resulted in juice delivery. Monkey U was trained in a task in which a center target appeared, requiring the monkey to hold it for 1 second before a second target appeared on the left or the right of the screen. The monkey had to release the holding target within 1 second to receive the juice reward. We first aligned the calcium transients to the appearance of the rewarded target, as an approximation of movement onset.

We found that in monkey Q’s SMA, 34% (43/129) of cells significantly modulated their activity in response to the presentation of the rewarded target and the immediately ensuing movement (p < 0.05, Wilcoxon signed rank test, corrected for multiple comparisons), as shown in figure 2A, where the average raw calcium traces of 129 cells (obtained across 7 sessions) was aligned to target presentation. In 11/129 cells (9%), the activity was not direction-related, but 32/129 cells (25%), showed larger changes in activity in trials where the target was located at a specific position (figure 2B, 2C). The magnitude of the responses did not differ between cells responding with an increase or decrease in firing (Mann-Whitney rank sum test) and was similar regardless of the target position (Kruskal-Wallis test; data not shown).

**Figure 2:**
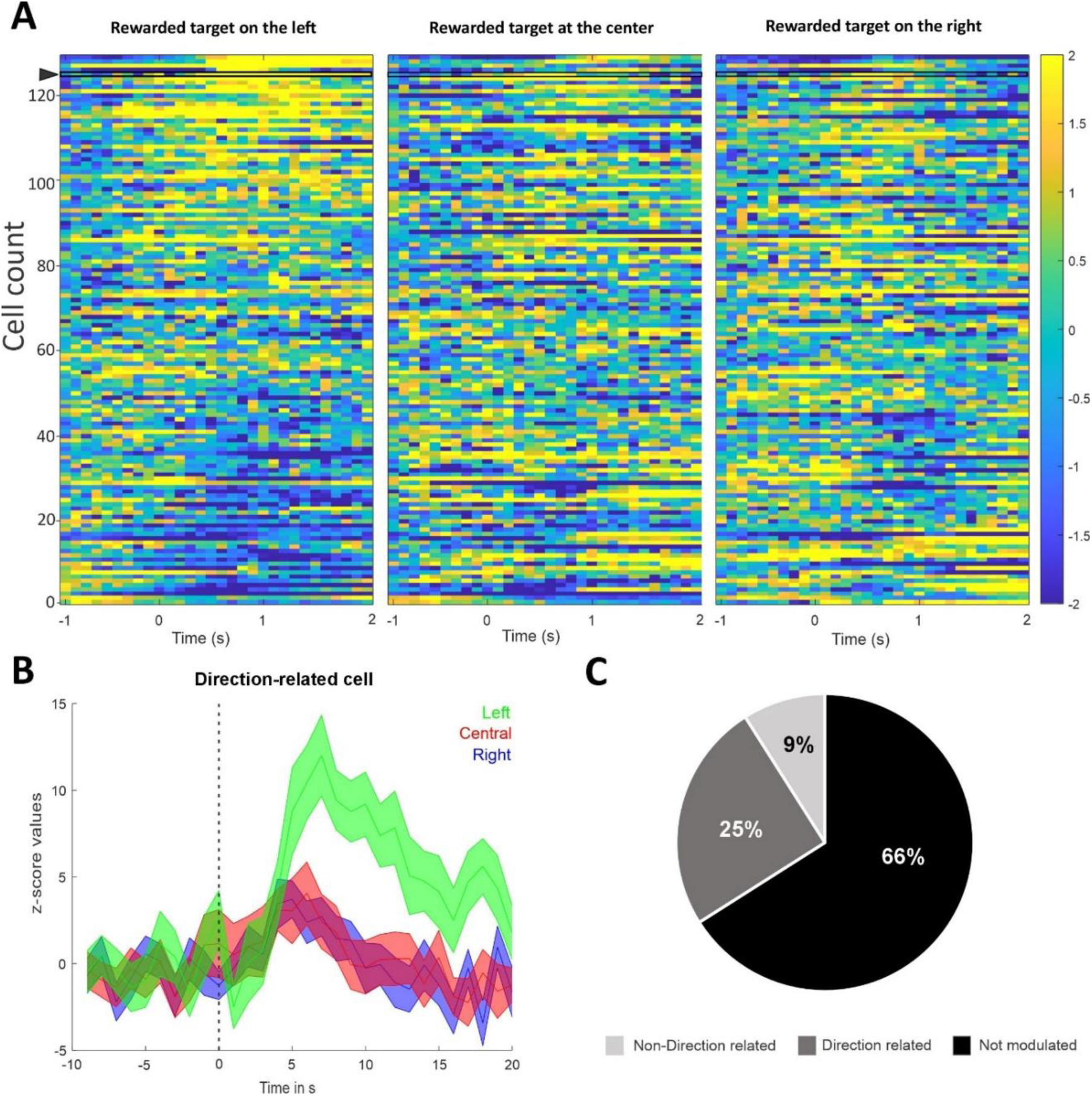
Calcium activity changes in SMA neurons related to the arm reaching task. **A:** Heat maps of the Z-scored raw calcium traces for each cell in the population, aligned on rewarded target onset on the right, center or left of the screen during the single-target reaching task in monkey Q (n=129 cells, average of 30 trials/conditions). In the left panel, the cells have been sorted, based on the amplitude of change in the Z-scored data. On the middle and right panel, cells are sorted in the same order as left panel. **B:** Example of a direction-related cell, indicated by the arrow in A, with a significantly higher increase of activity on the rewarded target’s presentation on the left. The cell’s activity is aligned on the rewarded target onset marked by the vertical dash line. Colored curves represent the average Z-score activity ± SD, separately for rewarded target on the left (green, p=0.03), central (red p=0.92) and right (blue, p=0.43). **C:** Pie chart representing the proportion of cells that were not modulated, direction-related or non-direction related (black, dark grey light grey respectively). Significance evaluated with an FDR-corrected Wilcoxon signed rank test with p<0.05.

In monkey U’s SMA, we found that 43% (6/14) cells had significant modulation of activity when the calcium traces were aligned to the appearance of the rewarded target, and in all cases (6/6) these changes in activity were related to the direction of the movement. These proportions were the same when the data was aligned to the movement onset (when the monkey removed the hand from the holding target).

In monkey U’s M1, when the calcium traces were aligned to the appearance of the rewarded target, 4/6 cells (67%) showed changes in activity, without directional relation. When the data was aligned to the movement onset, 100% (6/6) cells were modulated, and 50% of them showed direction-related activity (figure S1).

These data indicate that the calcium imaging method allows identification of a subset of SMA and M1 neurons that are modulated in a direction related manner when presented with a movement-related target in the reaching task.

### Cells in SMA and M1 show coactivation

We studied the synchrony between pairs of cells using the Jaccard index, using denoised data. Example sessions are shown for Monkey U in the SMA (figure 3A) and M1 (figure 3B) for spontaneous and task conditions. Pairs of neurons exhibited neural synchrony in both the SMA and M1 during the spontaneous and task conditions (figure 3A-B). The cell pairs exhibiting synchrony were not related to the centroid distances in either SMA or M1 (correlation plot in figure 3A-B). Figure 3C summarizes the proportion of synchronized pairs of cells (|Z-Jaccard| > 1.96) in the spontaneous and task condition recordings in SMA and M1 for all animals. The proportion of cells that were synchronized did not significantly differ between spontaneous and reaching tasks in either SMA or M1 (p=0.07 and p=0.22, respectively), however the number of sessions could be too low to produce significant differences. Cell-to-cell synchrony appears to be a reliable feature of SMA and M1 neurons, regardless of task and distance.

**Figure 3:**
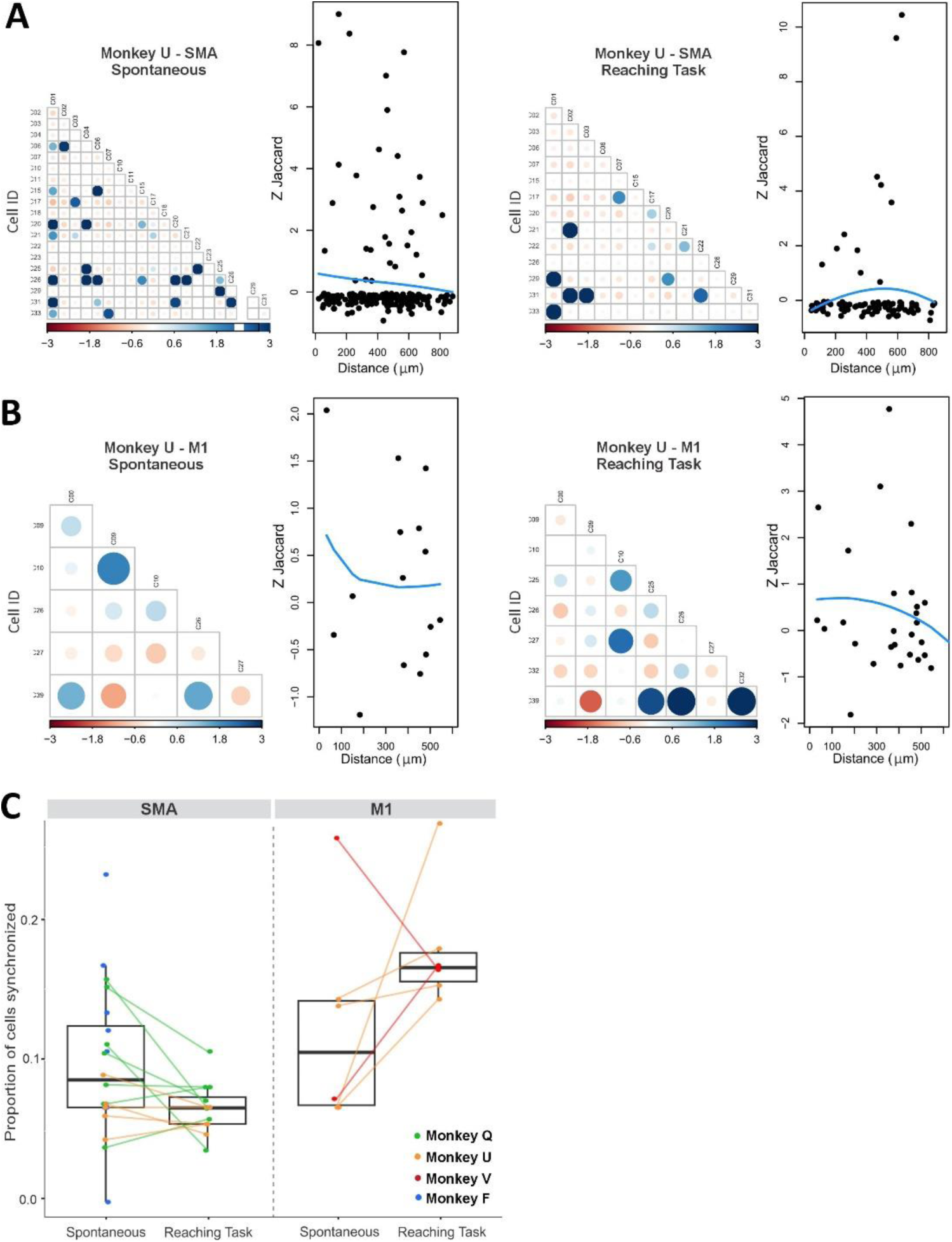
Coactivation between cells events in both SMA and M1. **A:** Coactivation plots based on recordings made during the spontaneous and the task conditions in SMA in one example session from monkey U. Circles represent normalized Jaccard index (Z-Jaccard) for each cell pairing. Large/dark blue circles indicate positive values of Z-Jaccard and large/dark red circles indicate negative values of Z-Jaccard. On the right of each coactivation plot, we show the correlation plots between Z-Jaccard index and distance between the centroids of the cells. **B:** Example session from monkey U in M1. Same legend as panel A. **C:** Box plot summarizing the proportion of cells synchronized during spontaneous and task conditions in SMA and M1. The horizontal bars represent the median value across all animals (monkeys Q, U, V and F in green, orange, red, and blue respectively). Each dot represents the average value for one session. Spontaneous and task values from the same session are connected by a line.

### Cells in SMA and M1 show sequential activation patterns

We then examined whether groups of neurons exhibited sequential activation patterns. A sequence was defined as a recurring series of calcium events involving at least 2 cells, occurring in a specific temporal order, and appearing at least 4 times withing a 10-minute recording. Sequences were only considered if the random occurrence probability was below 5% and if individual events were at least 0.05 seconds apart (to avoid detecting simultaneous bursts of events as ‘sequences’).

In an example imaging session in monkey Q’s SMA, we detected 13 such sequences in spontaneous and 17 sequences during the task (figure 4A, top panels). While some of the sequences detected involved the same cells (figure 4A bottom panels), the specific sequences differed (examples highlighted in green in figure 4A, 4B). We also performed the same analysis for sessions in which 20 minutes of spontaneous recording were collected, allowing us to segment the spontaneous segment in two 10-min periods (‘spontaneous 1’, ‘spontaneous 2’). An example from such a recording in monkey Q’s SMA is shown in figure 4B. We detected 8, 13 and 9 unique sequences during ‘spontaneous 1’, ‘spontaneous 2’ and the reaching task, respectively. The detected sequences differed between all three states, suggesting that the differences between the analyzed segments were not (solely) driven by the behavioral state.

**Figure 4:**
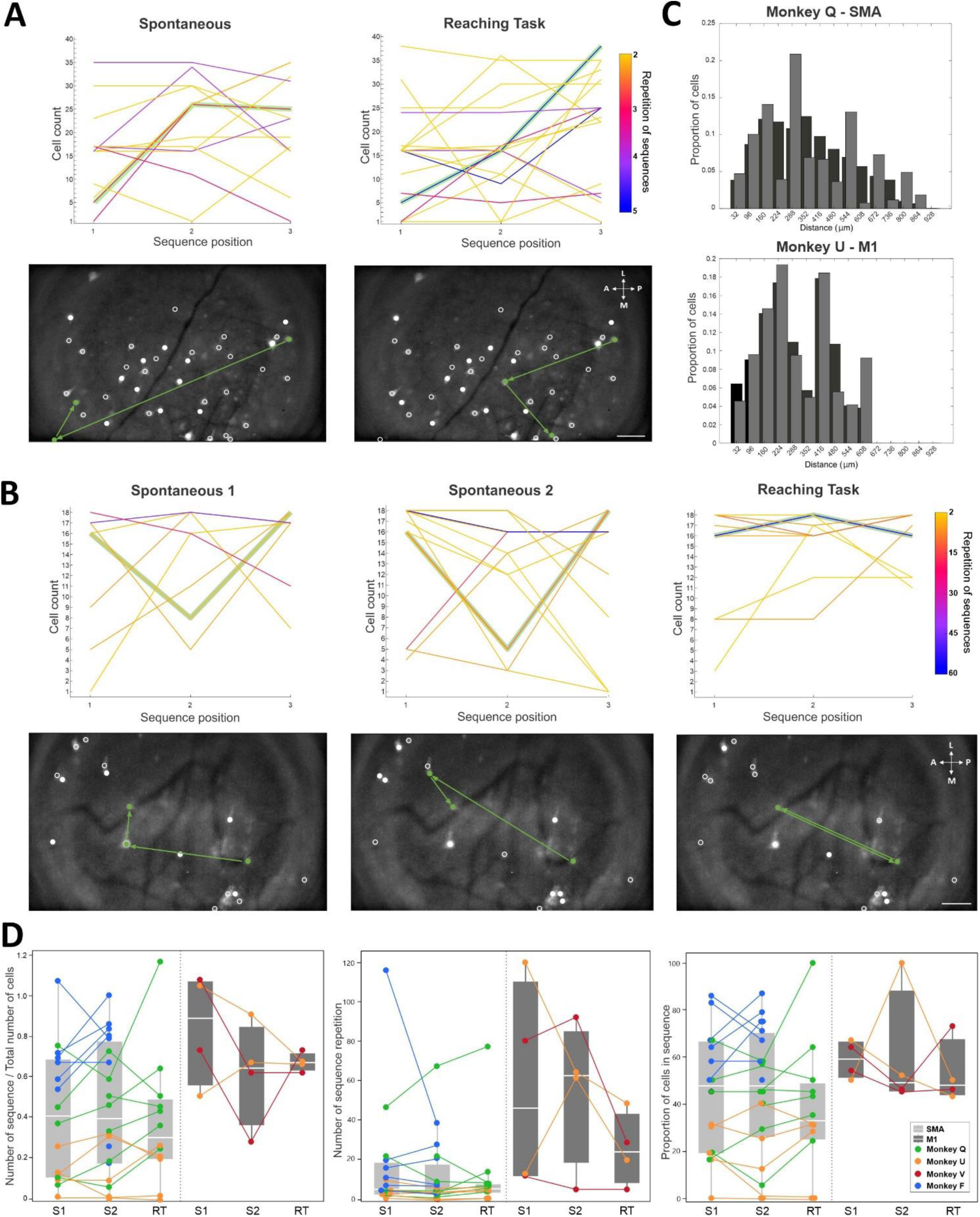
Repeating sequences of calcium events involving several neurons in the spontaneous condition and during the reaching task. **A:** Top row, depiction of sequences found across 40 cells in an example session in monkey Q’s SMA, during spontaneous and reaching task conditions. The x-axis indicates the position of the cell within a sequence. The line thickness indicates the number of repetitions of the sequences. An example sequence starting with cell #5 is highlighted in green for the spontaneous and for the task condition. Below each graph we show the maximal GCaMP6f fluorescence intensity overlapped with the map of cells detected (white circles). The green circles illustrate the three cells involved in the sequence and the green arrows indicate the temporal order. Although cell #5 initiates the sequence in both examples, it is followed by different cells in the spontaneous and reaching task. Scale bar: 100 µm. **B:** Sequences found across 18 cells in a different session in monkey Q’s SMA, during two 10 min segments of the spontaneous conditions (‘spontaneous 1’, ‘spontaneous 2’) and during the reaching task. Same conventions as described for figure 4A. **C:** Spatial distribution of cells involved in sequences. Black bars indicate the distribution of the centroids of all the cells recorded across all sessions in the spontaneous condition in monkey Q SMA (top) and monkey U M1 (bottom). Grey bars indicate the distribution of the centroids of the cells involved in sequences. Note overlap of the distributions, indicating that cells involved in sequences were not clustered, but spread across the field of view. **D:** Box plots representing normalized number of sequences, number of repetition and proportion of cell in sequence across monkeys in SMA (light grey) and M1 (dark grey). Each dot represents median values for a session. Monkey Q (green), U (yellow) F (blue) and V (red). Data is binned in 10 min of recording in spontaneous (S1 and S2) and during the reaching task (RT).

For all imaging sites, and for each recording segment, we calculated the number of sequences (normalized by the number of identified cells), the repetition rate of sequences and the proportion of cells involved in sequences (summarized in figure 4D). We found that these factors varied across conditions and across imaging sites, similar to observations made in monkey Q’s SMA.

Finally, we investigated whether cells participating in a sequence were spatially close. Plotting the distribution of distances between pairs of cells that were involved in sequences, as well as the overall distribution of distances between all detected cells showed that there is no preferred spatial relation between cells involved in sequences (figure 4C and figure S2).

### Expression of GCaMP6f and lens localization

Post-mortem histological examination confirmed that the lens and prism were placed in the arm regions of SMA and M1 ^8,13–15^ (figure S3A). However, in monkey Q, the SMA lens was positioned at the anterior edge of this region, likely bordering with the pre-SMA area ^8,15^.

The reconstruction of the location of the lens/prism showed that the probes were targeted to the deep cortical layers of the SMA/M1 (right panels in figure S3A). GCaMP6f expression around the imaging sites was revealed by immunoperoxidase against GFP (figure S3B) or by studying the endogenous fluorescence of the calcium sensor (figure S3C). We observed robust GFP expression around the lens/prism for most sites, except in monkey U’s SMA, where the GFP expression was sparse. Importantly, many GFP-positive cells showed the morphology of pyramidal cells (i.e., excitatory projection neurons) ^16^. We confirmed that GCaMP6f expression was specific for non-GABAergic neurons by performing a double immunofluorescence assay for GABA and GFP (figure S3D), confirming that GCaMP6f-positivecells are likely to be projection neurons and not interneurons.

## DISCUSSION

We examined the activity of projection neurons in deep layers of SMA and M1 of four rhesus macaques, when they were at rest (spontaneous condition) or performing a simple arm reaching task, using calcium imaging techniques. As expected, based on the results of previous electrophysiologic studies, we found that a proportion of SMA and M1 neurons generated calcium transients in relation to arm reaches, and that these responses were often directionally-related. We also identified pairs of projection neurons in SMA and M1 with coincident activity, and dynamically changing sequential activity patterns. Neither coincident nor sequence-related activities differed by spatial distances within the imaged field.

Our study complements the extensive literature on the electrophysiological activity of SMA and M1 neurons in macaques ^2^, and it builds upon previous studies that have used calcium imaging and miniscopes to examine motor cortex activity in rodents (e.g., ^17,18^). The calcium imaging method offers unique opportunities to increase our understanding of cortical activity in NHPs, allowing us to image many neurons in deep layers in M1 and SMA. As an important distinction to the previous electrophysiologic studies in rhesus monkeys, by combining a genetic strategy (use of the Thy1 promoter which targets preferentially projection neurons ^12^) and placing the GRIN lenses in deep cortical layers, we selectively monitored cortico-fugal neurons.

We compared calcium activity from the spontaneous to the reaching task condition in individually identified neurons that were reliably tracked within a session. We found that neither the rate nor the variability of calcium events differed significantly between the two conditions, perhaps indicating that the information coded by these parameters is not indicative of the behavioral state of the animal. However, a previous microendoscopic calcium imaging study in the M1 of marmosets reported an increase in calcium event rates from rest to movement ^19^ a difference that could be due to the more complex tasks the marmosets were engaged in (reaching for food pellets or climbing ladders).

We found that calcium event rates were, in general, higher in M1 neurons than in SMA neurons, which differs from electrophysiology studies which report similar firing rates in both regions ^15^. The difference may be partially explained by the fact that the electrophysiologic measurements were not specific for corticofugal neurons. In addition, the two techniques detect different neuronal activity patterns: while electrophysiologic recordings identify both single spikes and bursts of firing, calcium imaging is more sensitive to bursts of activity ^20^. Our result may therefore reflect a difference in resting firing patterns between M1 and SMA rather than a different overall activity of neurons in these regions.

In our study, the calcium event rate was about 10-fold higher than previously reported in the premotor cortex of rhesus macaques ^9^. The rate of calcium events we detected was also different from that reported in the M1 of marmosets ^19^. These discrepancies are likely not only related to species differences (in the case of the marmosets), but also a direct result of the definition of ‘calcium events’: our calculation of the event rate is based on the detection of such events with a deconvolution method, unlike the study of Bollimunta et al. (2021).

In a subpopulation of SMA and M1 neurons the calcium activity was related to arm reaches, and some neurons showed directional sensitivity, as previously observed with electrophysiological recordings ^2,8,15^.

However, the proportions of SMA neurons (34% for Monkey Q and 43% for Monkey U) showing task-related responses was lower to previous electrophysiological reports where monkeys engaged in similar arm-reaching tasks (e.g., refs. ^15,21–23^). This may have resulted from the placement of the lens in the most anterior portion of the SMA forelimb region, which has a patchier representation of the forelimb than the most caudal SMA ^8,15^. On the other hand, the proportion of neurons that showed task-related responses in M1 (100% when aligned on movement onset) aligns with findings from previous electrophysiological studies ^15,23^, even though the total number of neurons imaged in this location was low.

Similar results were reported in another study using microscopic calcium imaging in the M1 of marmosets ^19^. Thus, even though the time resolution of calcium transients is orders of magnitude lower than electrophysiological recordings, the calcium imaging method can detect basic functional parameters of these cortical regions.

Microendoscopic calcium imaging can simultaneously monitor many cells in a relatively large area of cortex, allowing analysis of simultaneous and sequential activation of neurons. In SMA and M1 we found pairs of cells that were coactive, unrelated to their spatial proximity (within the confines of the evaluated imaging field). Our results in the macaque motor cortices differ from the findings of Parker et al, who reported coactivation in spatially close groups of striatal medium spiny neurons in rodents ^24^, likely due to different structural organization across brain regions.

In a related analysis, we found that the activity of neurons in M1 and SMA can form recurring sequential patterns, involving multiple cells. Similarly to the findings of the coactivation analysis, the distance between neurons did not predict their involvement in sequences. The sequences were not stable over time, and not specifically dependent on the behavioral context of the recordings, as variations of patterns were also seen when comparing different segments of recordings done in the spontaneous condition. We could not derive a true estimate of the variability of the statistically identified sequences because the low calcium event rates preclude an examination of these sequences in short time intervals. Given that the identified sequence interactions occurred over relatively long periods (up to 2 seconds), they are not likely to be the product of local neuronal interactions but may represent long-range network activities.

As far as we know, this is the first report of the use of calcium imaging to examine neuronal sequential activation patterns in rhesus macaques. Sequential activation patterns of neurons have been studied before, e.g., in electrophysiologic and calcium imaging studies of bird song (e.g., ^25,26^) where sequential patterns can be traced to precise vocal patterns. As mentioned, we have not found clear links of sequence patterns to behavior in our animals, but it is possible that the detected sequences could be related to functional processes that were not directly assessed in our study, such as attentional or motivational states.

The calcium imaging technique has some important limitations. The temporal resolution of the calcium transient imaging procedure is much lower than that of electrophysiological recordings, limiting the analysis of neuronal events that occur at faster time scales. One reported advantage of the calcium imaging technique using miniscopes is the ability to track the same cells across days ^27^, based on the spatial stability of fluorescence and surrounding tissue landmarks within the imaged field. However, in our study we were not able to routinely track neurons across multiple sessions. This issue was also reported in studies conducting calcium imaging in the M1 of rats ^28^. Several factors may impact the ability to longitudinally track calcium transients in the same cells. One factor could be that the expression of GCaMp6f is unstable, but this seems unlikely, as our postmortem studies showed robust calcium sensor expression even after survival times of up to 189 days following viral vector injections. Alternatively, changes in the field of view, due to movement of the brain tissue in relation to the GRIN lenses, may interfere with longitudinal recordings. It is conceivable that this may differ between animal species, or even between brain regions within the same animal.

The sparse GCaMp6f expression in monkey U’s SMA may explain the low number of cells imaged in this animal. However, the GCaMP6f expression levels observed post-mortem in monkeys F and V, and in monkey U’s M1 were much higher, but the number of cells imaged *in vivo* were also found to be comparatively low. Overall, the total number of neurons imaged per session in our studies ranged from 23 to 129 (Table S4), which contrasts to a reported average of 90 cells in the rat M1 ^28^. This difference may reflect the fact that the neuron density is lower in the primate M1 compared to that of the rodent M1 ^29,30^.

In summary, we used microendoscopic calcium imaging to study groups of neurons in the SMA and M1 in rhesus macaques, across behavioral states. Our study sets the stage for use of this method to examine calcium dynamics in groups of cells in the primate motor cortices in more complex and controlled behavioral tasks, to compare patterns of activity among cells in disease states, such as parkinsonism ^31^, and to study neuronal responses to acute or chronic treatments in these models. Finally, given the ongoing development of enhancer sequences that can be used in AAVs or other viral vectors ^32^, it may soon be possible to expand this technique to track the activity of other neuronal subtypes (e.g., subtypes of cortical inhibitory interneurons, or specific cortical projection neurons).

## STAR METHODS

Detailed methods are provided in the online version of this paper and include the following:

- Key resources table
- Resource availability

- Lead contact
- Material availability
- Data and code availability
- Experimental model and study participant details
- Method details

- Behavioral task
- Surgery (AAV injections and lens implant)
- Calcium imaging data acquisition
- Histology
- Quantification and statistical analysis

- Cell identification and trace extraction
- Statistical comparisons of calcium activity rates and amplitudes
- Alignment to behavioral events
- Cell coactivation (Jaccard index method)
- Determination of precisely timed sequences of calcium events

## ACKNOWLEDGMENTS AND FUNDING SOURCES

We are grateful to Wendy Williamson Coyne for assistance during surgeries, and to Susan Jenkins, DeErra Locklin and Charlotte Armstrong for technical help. Some of the surgeries described in this study were conducted in collaboration with the Emory Translational Neuroscience Core, which is supported by the Department of Neurosurgery, Emory University School of Medicine.

This research was funded by Aligning Science Across Parkinson’s [ASAP-020572] through the Michael J. Fox Foundation for Parkinson’s Research (MJFF) and by NIH P51-OD011132 (Emory National Primate Research Center).

## AUTHORS CONTRIBUTIONS

Conceptualization, J.J.N, T.W., A.G.; Methodology, J.J.N., W.Y., J. D., T.W., A.G.; Software, B.R., A.D., T.W.; Investigation, A.C.M, D.P., A.G.; Formal Analysis, A.C.M., D.P., A.D., B.R., T.W.; Writing – Original Draft, A.C.M, D.P., A.G.; Writing – Review & Editing, A.D., B.R., J.J.N., W.Y., T.W., Visualization, A.C.M., D.P., A.D., B.R., T.W., A.G.; Supervision, T.W., A.G., Funding Acquisition, T.W., A.G.

## STAR METHODS

### KEY RESOURCES TABLE

| REAGENT OR RESOURCE | SOURCE | IDENTIFIER |
| --- | --- | --- |
| **Plasmids** |  |  |
| pAAV TREGCAMP6f | Addgene | Cat # 97410 |
| pAAV ThyStTA | Addgene | Cat # 97411 |
| **Viral vectors** |  |  |
| AAV5-TRE-GCAMP6f | Vigene Biosciences | n/a |
| AAV5-ThyStTA | Vigene Biosciences | n/a |
| **Antibodies** |  |  |
| Chicken anti-GFP (immunofluorescence) | Millipore | Cat # 06-896 RRID: AB_310288 |
| Rabbit anti-GFP (immunoperoxidase) | Thermo Fisher (Invitrogen) | Cat # A-11122 RRID: AB_221569 |
| Mouse anti-MAP2 | Millipore | Cat # MAB3418 RRID: AB_11212326 |
| Rabbit anti-GABA | Sigma | Cat # A2052 RRID: AB_477652 |
| Biotinylated goat anti-rabbit | Vector | Cat # BA1000 RRID: AB_2313606 |
| Biotinylated goat anti-mouse | Vector | Cat # BA9200 RRID: AB_2336171 |
| Donkey anti-chicken (Fluorescein) | Jackson Lab | Cat # 703-095-155 RRID:AB_2340356 |
| Donkey anti-rabbit (Rhodamine red-X) | Jackson Lab | Cat # 711-295-152 RRID:AB_2340613 |
| **EXPERIMENTAL MODELS: ORGANISM/STRAINS** |  |  |
| Rhesus macaques (*Macaca mulatta*) | Emory National Primate Research Center, Atlanta, GA, USA. | N/A |
| **PROTOCOLS** |  |  |
| Surgery to inject viral vectors and implant GRIN lenses | This study | [dx.doi.org/10.17504/protocols.io.e6nvw15w2lmk/v1](https://doi.org/10.17504/protocols.io.e6nvw15w2lmk/v1) |
| Miniscope calcium imaging data acquisition of cortical activity | This study | [dx.doi.org/10.17504/protocols.io.14egn612pl5d/v1](https://doi.org/10.17504/protocols.io.14egn612pl5d/v1) |
| Arm reaching to touchscreen behavioral task | This study |  |
| Immunoperoxidase staining | This study | [dx.doi.org/10.17504/protocols.io.14egn2b6mg5d/v1](https://doi.org/10.17504/protocols.io.14egn2b6mg5d/v1) |
| **SOFTWARE AND ALGORITHMS** |  |  |
| IDPS | Inscopix | https://inscopix.com/software-analysis-miniscope-imaging/ |
| Matlab | MathWorks | https://www.mathworks.com/ |
| R | R Core Team | https://www.r-project.org/ |
| CorelDraw |  | https://www.coreldraw.com/ |
| **DEPOSITED DATA** |  |  |
| Raw data and Code | This study | Pending* |
| *Items pending will be provided if accepted for publication |  |  |

### RESOURCES AVAILABILITY

#### Lead contact

Further information and requests for resources should be directed to and will be fulfilled by the lead contact, Anne-Caroline Martel.

#### Material availability

This study did not generate new materials.

#### Data and code availability

- Data: All data generated in the study have been deposited at DANDI and will be made publicly accessible if the paper is accepted for publication. The corresponding DOI will also be listed in the key resources table.
- Code: All original code used in the study have been deposited at Zenodo and will be made publicly accessible if the paper is accepted for publication. The corresponding DOI will also be listed in the key resources table.
- Any additional information required to reanalyze the data reported in this paper is available from the lead contact upon request.

### EXPERIMENTAL MODEL AND SUBJECT DETAILS

Four rhesus macaques (Macaca mulatta) were studied (monkeys F, Q, U, and V, 2 males, 2 females). The animals were 4 - 5 years old and weighed 5.5 - 6.8 kg at the beginning of the study. The monkeys were pair-housed in a temperature-controlled room with a 12-hour light cycle, fed a standard primate diet twice daily with additional fruit and vegetable supplements, and had access to water *ad libitum*. All procedures were approved by the Institutional Animal Care and Use Committee (IACUC) of Emory University and were performed according to the Guide for the Care and Use of Laboratory Animals and the U.S. Public Health Service Policy on the Humane Care and Use of Laboratory Animals.

### METHODS DETAILS

#### Behavioral task

Two of the monkeys (Q and U) were first trained to accept head fixation with a thermoplastic molded helmet (CIVCO medical solutions) (Drucker et al., 2015) or a surgically implanted head post, and then trained on an arm reaching task (NIMH, MonkeyLogic v2.2). The monkeys were facing a 17-inch display touchscreen (GVision) during task performance. Training began 2-4 weeks before lens implant surgery (see below) and continued until the animals reached criteria (see below).

Monkey Q was trained in a one-target task that required the animal to touch a circle (∼93mm diameter) shown on the touchscreen display with its right hand to receive a juice reward. For each trial, the target would randomly appear on the right, left, or center of the touchscreen and remained on display for up to 3s. If the monkey touched it during this period (successful trials), the target disappeared, and the animal received a drop of juice. A new trial started after a random inter-trial interval (2-3 seconds), following the release of the monkey’s hand from the touchscreen.

Monkey U was trained in a two-target task, in which the monkey had to first touch a center target (circle, ∼52 mm diameter). After the monkey held the center target for 1 second, targets on the left or right side of the screen (randomly chosen) would appear. The monkey had to release the center target within 1 second and touch the side target (successful trial), to receive a drop of juice. The monkey worked with its right hand for the imaging sessions evaluating the left SMA, or with his left hand for the imaging sessions focused on the right M1.

We considered monkeys trained in these tasks when they consistently completed at least 100 successful trials per 10-minute session.

#### Surgery (AAV injections and lens implant)

Monkeys were subjected to a surgery to receive injection of virus solutions and implantation of GRIN lenses, following previously described methods (see “Key Resources Table” and Bollimunta et al.(2021)). The location of the lens implants was determined based on pre-surgery MRI scans and stereotaxic coordinates obtained from (Paxinos et al., 2000). The surgery was performed under sterile conditions. Monkeys were sedated with ketamine (10mg/kg), followed by isoflurane anesthesia (1-3%) for the remainder of the procedure. They were placed in a stereotactic frame, and their vital signs were monitored throughout the surgical procedure.

We performed two craniotomies, each producing skull defects measuring approximately 10 mm in diameter, over the arm regions of the left SMA and the right M1. Following the craniotomies, we made a small incision in the dura mater for virus injections and GRIN lens implants at each cortical site.

We used a virus solution consisting of a 1:1 mixture of AAV5-TRE-GCAMP6f (9.05×10e13 gc/ml) and AAV5-ThyStTA (5.13×10e13 gc/ml). The Tet-Off system strategy was used because the GCaMP6f expression is amplified due to the tetracycline (tTA) promoter activation of the tetracycline response element (TRE3) (Bollimunta et al., 2021; Kondo et al., 2018). In addition, if needed, GCaMP6f expression could be temporarily suppressed with systemic administration of doxycycline (Bollimunta et al., 2021). Note, however, that doxycycline was not used in these studies. A Hamilton syringe with a 25-ga needle was filled with the mixture and then lowered into the brain using an injection pump mounted to the stereotactic frame (Stoelting quintessential stereotaxic injector). We started injections of the virus solution after a three-minute wait time.

We injected 3-6.4 and 5-7.8 µl (at 0.1 – 0.8μl/min) of virus solution in M1 and SMA, respectively. Information about the depths and volumes of injections in each site are provided in supplementary material (Table S2). The needle remained at each injection site for at least 3 min after the final injection.

After the virus injection, an epoxy-filled 18-ga needle was lowered into the brain to create a path for the subsequent insertion of a GRIN prism for M1 (1 mm diameter, 9 mm, or 12 mm length), or a flat lens for SMA (1 mm diameter, 12 mm length), pre-attached to a baseplate to dock a miniscope (ProView Integrated Probe or ProView Integrated Prism Probe; Inscopix). In both brain regions we aimed to implant the lenses at a vertical or nearly vertical orientation (for ease and stability of later microscope docking) and image perpendicular to the plane of the cortical surface. For M1, which sits on the gyral surface of the cortex, a prism is required to attain the intended imaging field of view. For SMA, which sits in the cingulate sulcus along the medial wall of the brain, no prism is necessary and therefore we used a flat lens. The baseplate with the integrated lenses were lowered into the brain to target the center of the dorsal-ventral extent of the virus injections. Following the lens implantation, a cranial chamber was positioned around each baseplate-integrated lens, affixed to the skull, and capped for protection. The exposed brain tissue around the lens was covered with gel foam and a slotted aluminum disk. The implant was secured to the disk and skull within the chamber with Metabond dental cement (Parkell, Edgewood, NY) or Gradia Direct Flo light-cured composite (GC Corp., Tokyo Japan). Finally, the exposed skull was covered with bone screws and dental acrylic. We then secured a head holder in the dental acrylic cap.

#### Calcium imaging data acquisition

Nineteen to 48 days post-surgery, we started imaging calcium transients in SMA or M1 neurons using a head-mounted miniscope (nVista3.0, Inscopix, CA, USA). We did not perform simultaneous recordings in both regions. The miniscope was docked to the integrated baseplate of the implanted lenses. Imaging data for each site were acquired at least twice weekly. During these sessions, the monkeys were sitting awake on a primate chair (confirmed by continuous monitoring via a live video stream). Head movement was restricted by a thermoplastic helmet or by a head post. In each session, we performed 10-20 minutes of imaging in the ‘spontaneous’ condition during which the monkeys performed only occasional spontaneous movements of arms, trunk, and legs, followed by imaging while the animals worked on the ‘reaching task’ for as long as it took to complete 100 successful trials (median 12 min, inter-quartile range (IQR) 3.5 min). For some of these recordings, TTL pulses from the behavioral task, indicating the occurrence of behavioral events, were recorded on additional channels. The miniscope remained mounted until the end of the session. Imaging and recording parameters were controlled using the Inscopix Data Acquisition Software (IDAS, Inscopix; frame rate, 10 Hz; LED power, 0.6-0.8 mW/mm^2^; sensor gain, 7-8X; and electronic focus, 400-900).

#### Histology

Monkeys were euthanized at the conclusion of the experiments with an overdose of pentobarbital sodium (100mg/kg, iv). The post-surgery survival times were 189 days for monkey Q, 121 days for monkey U, 118 days for monkey V and 78 days for monkey F. They were then transcardially perfused with oxygenated Ringer’s solution, followed by 2 liters of fixative [4% paraformaldehyde, 0.1% glutaraldehyde in phosphate buffer (0.1 M, pH 7.2)]. The fixed brains were then removed from the skull and blocked. The blocks were later cut into 60 µm slices using a vibratome and immunostained for the neuronal marker microtubule-associated protein 2 (MAP2), and for green fluorescent protein (GFP) to examine lens placement and GCaMP6f expression in neuronal cell bodies of projection neurons, respectively. In addition, to confirm the identity of the GCaMP6f expressing neurons, we performed double fluorescent immunolabeling for GABA (as a marker of interneurons). See “Key Resources Table” for antibody details.

### QUANTIFICATION AND STATISTICAL ANALYSIS

#### Cell identification and trace extraction

Raw calcium imaging recordings, comprised of spontaneous and reaching task conditions, were imported into the IDPS (Inscopix Data Processing Software) for cell identification and trace extraction. The recordings were preprocessed by removing excess pixels, correcting defective pixels using a 3×3 median filter, and performing 4-fold spatial down sampling of each image frame. We then applied a spatial band-pass filter (low and high cut-off points of 0.005 pixel^-1^ and 0.5 pixel^-1^, respectively) and performed rigid translation motion correction with a mean projection image as the global reference frame (to which other frames were aligned). Following that, we used the constrained nonnegative matrix factorization for microendoscopic data (CNMFe) algorithm (Zhou et al., 2018) to extract the calcium transients of individual cells. From the extracted calcium traces, we identified and accepted cells based on imaging metrics such as the cell’s spatial footprint, size, and signal-to-noise ratio. Profiles that did not have a somatic appearance (e.g., blood vessels or dendritic segments) or calcium dynamics were rejected from further analysis. On accepted cells, we deconvolved the traces using the “Online Active Set methods for Spike Inference” (OASIS) method (Friedrich et al., 2017; Zhou et al., 2018) to obtain the calcium events per cell. For each recording, we calculated the distance between the cells based on their spatial centroids (X Y coordinates). All data were further analyzed in MATLAB R2023b and R (v 4.4.1). We only report results for the first session in which cells were identified.

#### Statistical comparisons of calcium activity rates and amplitudes

For each identified cell, we calculated calcium event rates, the coefficient of variation of the inter-event intervals (IEI CV), and the mean amplitude of events during spontaneous and/or reaching task recording conditions. Statistical comparisons, contrasting the calcium event rates in the spontaneous and reaching task conditions in the SMA or M1, were performed using the Wilcoxon signed rank test. We were also able to compare data recorded in M1 and SMA (in monkey U only) in either the spontaneous or task conditions. These comparisons were performed using Wilcoxon rank sum tests. Additionally, we corrected for multiple comparisons using Benjamini-Hochberg false discovery rate (FDR) (Benjamini and Hochberg, 1995). Statistical significance was set at *p* < 0.05.

#### Alignment to behavioral events

We used TTL pulses generated during the behavioral task to align raw calcium traces to behavioral events, using a custom-written MATLAB algorithm. We analyzed data from seven recording sessions in monkey Q’s SMA, as well as one session in SMA and one in M1 from monkey U.

Since each monkey performed a slightly different task (monkey Q performed a one-target task, while monkey U was trained in a two-target task), we first aligned the calcium traces to the event common to both tasks: the appearance of the rewarded target that prompted a reaching movement for a juice reward. We considered this event a suitable proxy for movement onset, as the monkeys initiated an arm movement to touch the target upon its appearance on the screen. For monkey Q, this target was the first and only one in each trial, whereas for monkey U, the rewarded target was the second one, following a required hold period at the center target (see Behavioral Task). For each successful trial, the 3-second epoch of raw data starting 1 second before target appearance was Z-scored to the first second of the 2 second inter trial period.

Additionally, for monkey U we aligned the calcium traces to the moment when the monkey removed its hand from the holding target and began reaching for the rewarded target. This event may have more precisely reflected movement onset. In this case, we also Z-scored a 3-second epoch of raw data to the inter trial period.

For both analysis (alignment to the appearance of the rewarded target and alignment to movement onset), the mean and standard deviation of Z-scores were then calculated for each cell and trial conditions (target on the right, center, or left for monkey Q, and target on the right or left for monkey U). Results from example sessions are shown in figure 2 and figure S1. Cellular activity was considered as being modulated by the task events if the mean calcium activity during the event window (0-1 second after the reward target appeared for both animals, or 0-1 seconds after the hand left the center target for monkey U) differed significantly from the mean activity during the second preceding the target presentation or movement onset. We used Wilcoxon signed rank tests with multiple comparisons corrected at the level of each chamber using FDR. In the SMA of monkey Q, 129 cells were tested across 3 conditions (387 tests); for monkey U, 14 cells were tested in 2 conditions (28 tests) in SMA; and 6 cells were tested in 2 conditions (12 tests) in M1. Cellular activity was considered related to movement direction if only one condition was significant at FDR-corrected p-value <0.05, while non-directionally related neurons were those with a significant response to two or more conditions. The absolute value of the amplitude of change in the Z-scored signal between the 1-second preceding the target presentation and the 1-second following the target presentation was also extracted. The magnitude of change in cells showing an increase in activity, and those showing a decrease were compared using a Mann-Whitney rank sum test and comparisons across condition (right, center, or left target) were made with a Kruskal-Wallis test. In all cases, significance was assumed if p < 0.05.

#### Cell coactivation (Jaccard index method)

We examined the synchrony between calcium events using the Jaccard index (implemented using custom R code). Using the deconvolved calcium event data exported from IDPS, we analyzed all sessions from each site (SMA and/or M1). Following (Parker et al., 2018), we first binned the deconvolved calcium events data into 0.2 second intervals, resulting in a number of bins ranging from 1508 to 6547 (median=6012) and 2371 to 6041 (median=3700) for the spontaneous and reaching task data respectively. Then, we forward-smoothed that data by setting a bin equal to one if a calcium event occurred at any time during the current bin or four bins ahead (a total of 1 second). We then calculated the Jaccard index between all cells (the intersection of the number of forward-smoothed events divided by the union). For each cell pairing, we generated the null distribution of Jaccard indices by randomly circularly shifting the data and recalculating the Jaccard Index 1,000 times. This was repeated 1,000 times. The circular shift preserves the autocorrelation structure of the forward smoothed event data. Next, for each cell pairing, we calculated a normalized version of the Jaccard index by subtracting the mean of the randomly shifted Jaccard indices from the observed Jaccard index and dividing by the standard deviation of the randomly shifted Jaccard indices, which we refer to as the Z-Jaccard. Large positive values of Z-Jaccard indicate cells pairs whose events co-occur substantially more than random circular shifts of the data. Large negative values indicate cell pairs whose events co-occur less than expected by chance. We defined the proportion of cells that were synchronized as the fraction of cell pairs in which the absolute value of the Z-Jaccard was greater than 1.96. For the sessions in which we had both spontaneous and reaching task condition, we compared whether there was a change in the proportion of cells synchronized using a Wilcoxon signed rank test.

We used the centroid information to examine whether there was a relationship between the Z-Jaccard and the distance between cells. A loess smoother (span=1) was used to visualize the relationship between Z-Jaccard and distance.

#### Determination of precisely timed sequences of calcium events

Sequences of calcium events were detected using a custom-written MATLAB algorithm. Sequences of events were defined as the occurrence of two or more calcium spike events (based on the deconvolved data) in a defined temporal order (e.g., calcium event in neuron A -> calcium event in neuron B -> calcium event in neuron C) among two or more cells, with each step of the sequence occurring within a specific time limit (2 seconds). We evaluated the occurrence rates of such sequences during 10 min segments of recordings during the spontaneous and arm-reaching task conditions. For this, the algorithm first tabulated the occurrence rates of all possible combinations of events within a given record. To test whether a specific combination of events occurred more frequently than would be expected by random alignment of the events, we generate 1,000 shuffled arrangements of the data, by shifting one of the data streams against the other, by random periods of time (using the MATLAB ‘circshift’ routine). This procedure (largely) retains the temporal order of events within the data streams but disrupts the potential temporal coupling between them. Sequences that were detected in the original data were accepted as being significant if they occurred more commonly than seen in 95% of the shuffled data (thus, were outside of the 95^th^ percentile of the shuffled data), if they were found at least 4 times during the 10 min observation period, and if individual occurrences were separated by at least 0.05 seconds (to avoid detecting simultaneous bursts of events as sequences). We then evaluated the number of ‘significant’ sequences, the number of steps forming a sequence, the number of sequence repetitions, and the spatial distance of neurons participating in the sequence members (using the centroid values, see above).

## SUPPLEMENTAL INFORMATION

**Table S1:** Parameters for cell identification and deconvolution.

| Preprocess |  |  | Spatial Filter |  | Motion Correction |  |  |
| --- | --- | --- | --- | --- | --- | --- | --- |
| Spatial Downsampling | Fix defective pixels (Yes/No) | Trim early frames (Yes/No) | Low | High | Global ref. frame | Use ROI (Yes/No) | Preserve input dimension (Yes/No) |
| 4 | Yes | Yes | 0.005 | 0.5 | Mean Image | Yes | Yes |
| CNMFe Cell Identification |  |  |  |  | OASIS Deconvolution |  |  |
| Avg. Cell Diameter (pixels) | Min. Pixel correlation | Min. Peak to Noise ratio | Gaussian filter width (pixels) | Merging threshold | Spike SNR Threshold |  |  |
| 7-20 | 0.75 - 0.80 | 8-10 | 8-20 | 0.3 | 2.5 |  |  |

**Table S2:** Number of sessions and cells identified per imaging site. *Indicates days during which GCaMp6F positive cells were imaged.

| Animal ID | Structure | Imaging period (days)* | Number of sessions | Cells identified per session (median (range)) | Total cells identified across sessions | Cells identified in more than one session |
| --- | --- | --- | --- | --- | --- | --- |
| Q | SMA | 147 | 7 | 18 (12 - 40) | 129 | 23 |
| F | SMA | 36 | 9 | 12 (1 - 16) | 61 | 17 |
| U | SMA | 50 | 5 | 16 (10 - 25) | 63 | 15 |
| U | M1 | 35 | 4 | 8 (6 - 21) | 37 | 6 |
| V | M1 | 21 | 2 | 12.5 (11 - 14) | 23 | 1 |

**Table S3:** Details of AAV injections.

| Site | No. of injection tracks | Number of deposits | Deposit dorso-ventral location (mm) | Volume delivered per deposit (μl) |
| --- | --- | --- | --- | --- |
| Monkey Q SMA | 1 | 3 | 7, 6.5, 5.5 | 2.3, 1.8, 1.4 |
| Monkey Q M1 | 1 | 2 | 4, 3 | 2, 1.3 |
| Monkey U SMA | 1 | 3 | 4.9, 4.4, 3.4 | 2, 1.5, 1.5 |
| Monkey U M1 | 1 | 2 | 3.7, 2.7 | 1.5, 1.5 |
| Monkey V SMA | 1 | 3 | 5.8, 5.3, 4.3 | 2, 1.5, 1.5 |
| Monkey V M1 | 1 | 2 | 5.3, 4.3 | 1.5, 1.5 |
| Monkey F SMA | 1 | 4 | 5.5, 5, 4.5, 4 | 1.6, 1.6, 3, 1.6 |
| Monkey F M1 | 1 | 4 | 6.7, 6.2, 5.2, 4.7 | 1.6, 1.6, 1.6, 1.6 |

**Figure S1:**
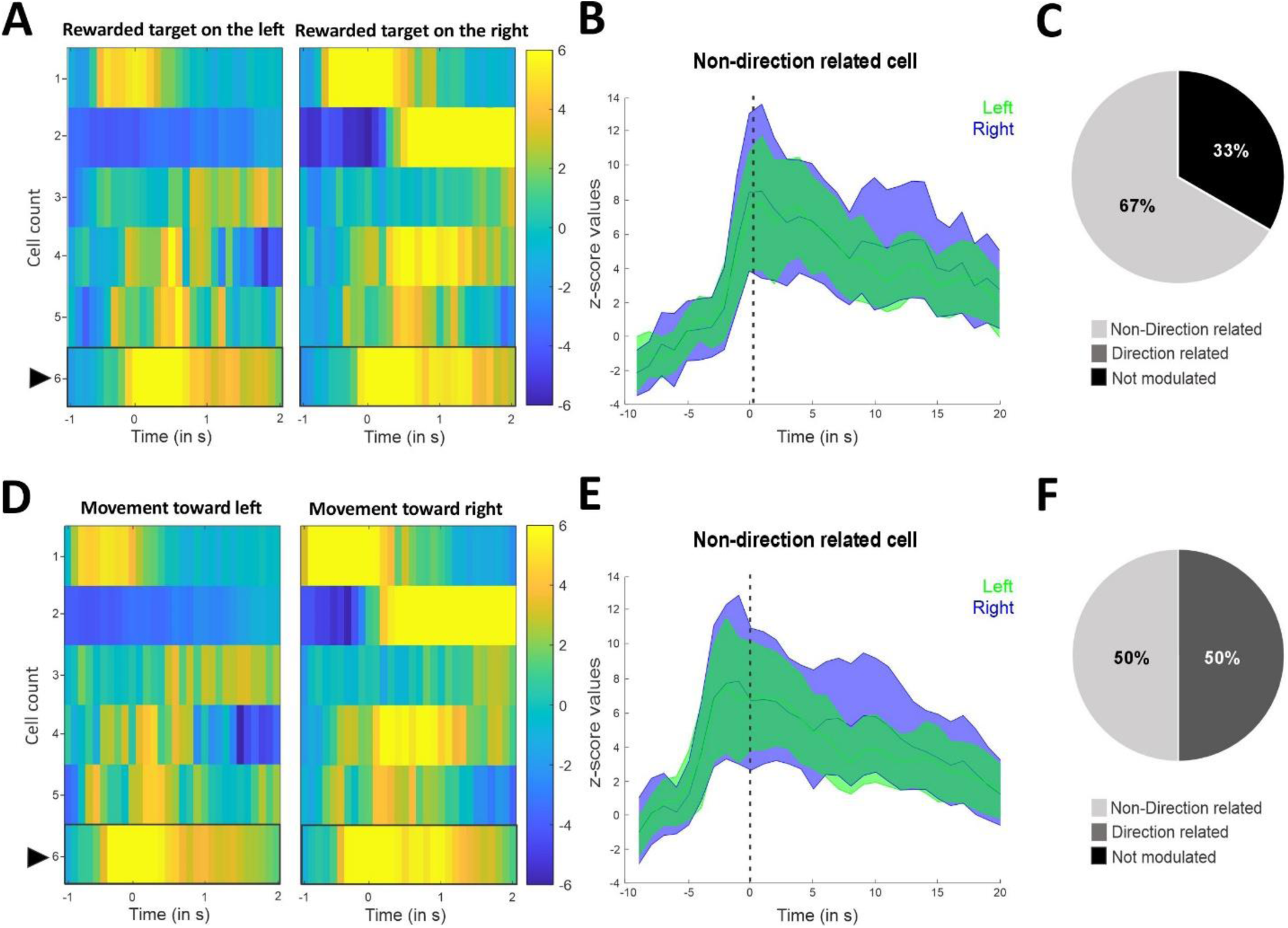
Changes in calcium activity in relation to rewarded target presentation and movement onset during an arm reaching task. **A:** Heat maps of the Z-scored raw calcium traces of each cell in an example session in M1, aligned on the rewarded target onset on the right or left during the two-target reaching task in monkey U (n=6 cells, 50 trials/condition). Each line indicates the same cell in both panels. **B:** Example of non-direction related cell, indicated by the arrow in A, with a significant increase toward both right and left rewarded targets (p<0.01 target left, p<0.01 target right, FDR-corrected Wilcoxon signed rank with p<0.05). The cell’s activity is aligned on the rewarded target onset marked by the vertical dash line. Colored curves represent the average z-score activity ± SD separately for left (green), and right (blue) targets. **C:** Pie chart representing the proportion of cells that are not modulated (black), direction related (dark grey, none in this case) and non-direction related (light grey). **D:** Heat maps of the Z-score raw calcium traces of each M1 cell in the same session shown in A-C, but aligned on the movement onset toward the right or left cue in monkey U. The cells are sorted in the same order as in panel A. **E:** Example of the same non-direction related cell presented in panel B (indicated by an arrow in D), aligned on movement onset, showing a significant increase in activity toward both right and left (p<0.05 target right, p<0.05 target left, FDR corrected Wilcoxon signed rank with p<0.05). F: Pie chart representing the proportion of cells that are not modulated (black, none in this case), direction related (dark grey) and non-direction related (light grey).

**Figure S2:**
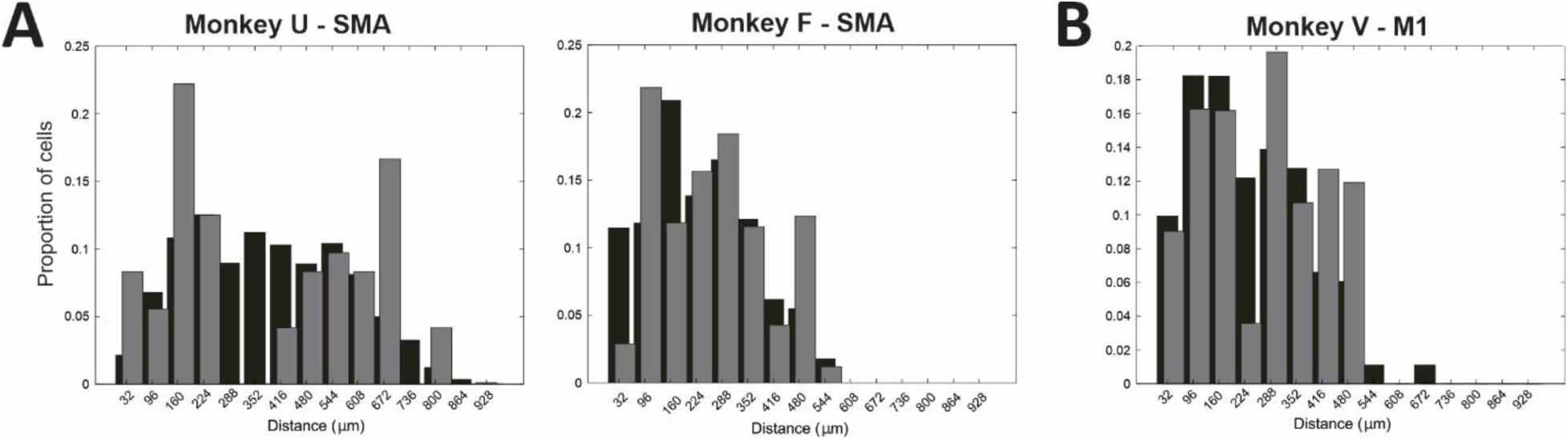
Spatial distribution of cells involved in sequences and sequence parameters. **A:** Black bars indicate the distribution of the centroids of all cells recorded across all sessions in the spontaneous condition for monkey U and F in SMA, monkey V in M1. Grey bars indicate the distribution of the centroids of the cells involved in sequences. Note almost complete overlap of the distributions, indicating that cells involved in sequences were not clustered, but spread across the field of view.

**Figure S3:**
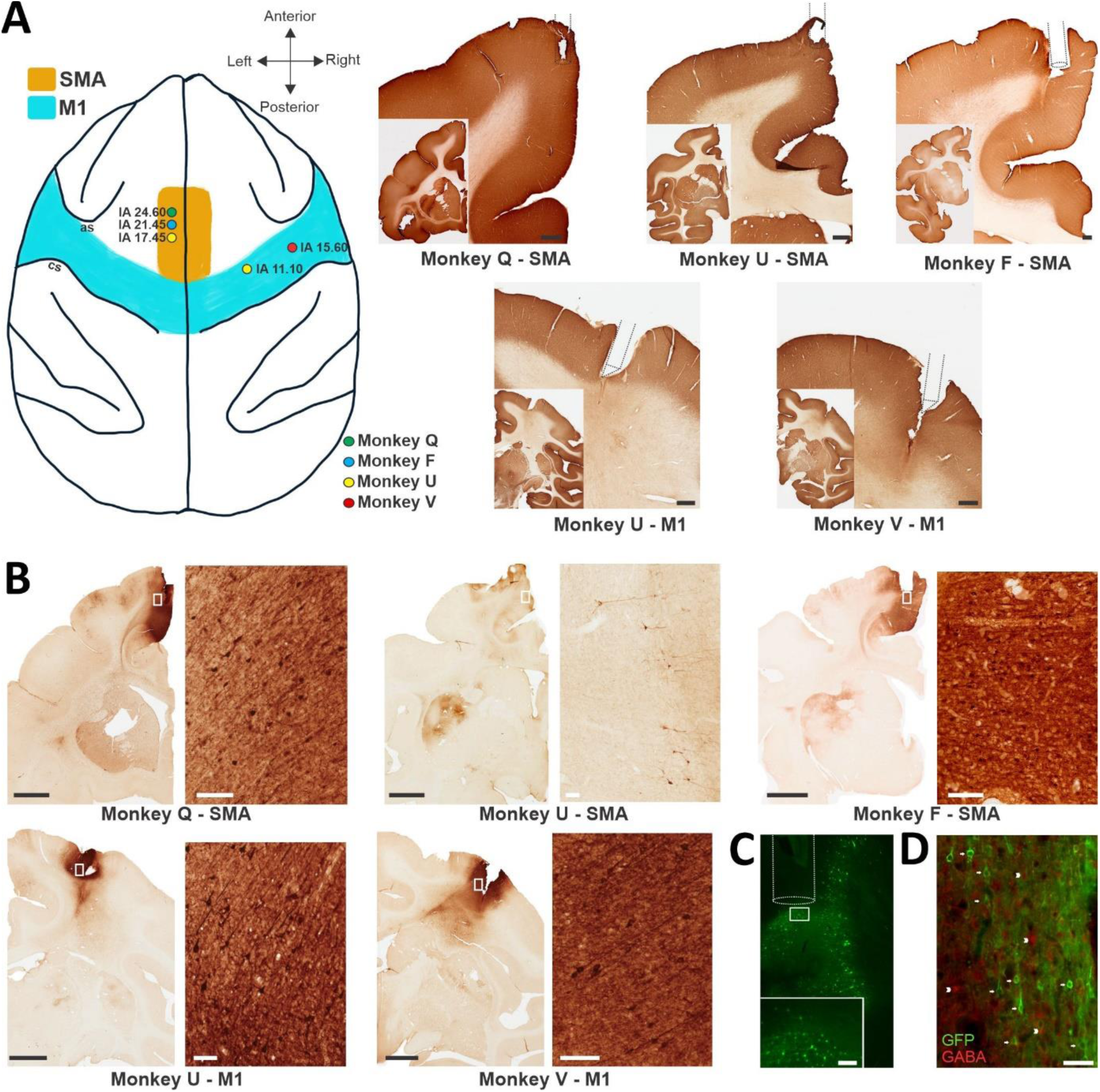
GCaMP6f expression and lens location in SMA and M1. **A:** This diagram indicates the location of GRIN lenses, based on post-mortem verification in SMA and M1 for monkeys Q (green dot), F (blue dot), U (yellow dots) and V (red dot) with the corresponding interaural coordinates (based on the atlas by Paxinos et al 2000). Abbreviations: as, arcuate sulcus; cs, central sulcus. Right: Reconstructions of the lens placement in SMA (top row) and prism placement in M1 (bottom row) in MAP-2 stained brain sections. Black scale bars, 1 mm. **B:** GFP immunoperoxidase (to reveal GCaMP6f expression) in SMA and and M1. The white rectangles are shown at higher magnification on the right. Black scale bars, 4 mm; white scale bars, 100 µm. **C:** Endogenous green fluorescence of GCaMP6f in a section adjacent to those shown in A and B for monkey Q. Reconstruction of lens placement (white line), lens diameter is 1 mm. The white rectangle is shown at higher higher magnification on the right. White scale bar, 100 µm. **D:** Double immunofluorescence for GFP and GABA in the M1 of monkey U. Green indicates GFP (GCaMp6f)-positive cells, some indicated by arrows). Red indicates GABA-positive cells (some indicated by arrowheads). No overlap of the signals was observed, suggesting that GCaMp6f-expressing cells are not GABAergic interneurons. White scale bar, 50 µm.

